# Enhanced and idiosyncratic neural representations of personally typical scenes

**DOI:** 10.1101/2024.07.31.605915

**Authors:** Gongting Wang, Lixiang Chen, Radoslaw Martin Cichy, Daniel Kaiser

**Affiliations:** Department of Education and Psychology, Freie Universität Berlin, Berlin, Germany; Department of Mathematics and Computer Science, Physics, Geography, Justus-Liebig-Universität Gießen, Gießen, Germany; Center for Mind, Brain and Behavior (CMBB), Justus Liebig University Gießen, Philipps-Universität Marburg and Technische Universität Darmstadt, Marburg, Germany

**Keywords:** Scene perception, Typicality, Individual differences, Drawing, EEG, Deep neural networks

## Abstract

Previous research shows that the typicality of visual scenes (i.e., if they are good examples of a category) determines how easily they can be perceived and represented in the brain. However, the unique visual diets individuals are exposed to across their lifetimes should sculpt very personal notions of typicality. Here, we thus investigated whether scenes that are more typical to individual observers are more accurately perceived and represented in the brain. We used drawings to enable participants to describe typical scenes (e.g., a kitchen) and converted these drawings into 3D renders. These renders were used as stimuli in a scene categorization task, during which we recorded EEG. In line with previous findings, categorization was most accurate for renders resembling the typical scene drawings of individual participants. Our EEG analyses reveal two critical insights on how these individual differences emerge on the neural level: First, personally typical scenes yielded enhanced neural representations from around 200 ms after onset. Second, personally typical scenes were represented in idiosyncratic ways, with reduced dependence on high-level visual features. We interpret these findings in a predictive processing framework, where individual differences in internal models of scene categories formed through experience shape visual analysis in idiosyncratic ways.

## Introduction

The ways in which humans perceive their environment are, by and large, studied across groups of participants, harnessing the coarse inter-individual stability of the mechanisms that guide visual perception. The focus on the group level is based on one of the core insights in vision research, which is that the visual system is remarkably similar across individuals and even across species (1-3). When taking a closer look, however, perceptual mechanisms differ across people: The cortical architecture of the visual system varies inter-individually, resulting in idiosyncratic biases in cortical processing (4-6). On the behavioral level, individual differences are observed in the perception of faces, objects, and expertise-related stimuli (7-10), as well as in the attentional allocation in faces, objects, and scenes (11-14).

One explanation for these idiosyncrasies lies in individually specific visual diets. The visual inputs we receive over our lifetimes (15, 16) sculpt the response properties on our visual systems in ways that give rise to individual perception and neural representation. Indeed, visual diets can be a powerful predictor for behavior and brain architecture (17, 18). If we take this argument seriously, and our visual representations are indeed formed by our unique visual diets, then what is a typical instance of a visual stimulus category, say a scene, is not necessarily typical for another individual – what is typical should vary as a function of what we learned about the scene category over our lifetimes.

Here, we test whether individual notions of typicality are linked to idiosyncrasies in visual processing, focusing on the perception of complex and naturalistic visual scenes. To quantify what constitutes a typical scene for each individual participant, we first asked our participants to draw typical instances of real-world scene categories (e.g., a kitchen or a bathroom), a paradigm we developed in recent behavioral work (19). We then conducted an EEG experiment, in which participants categorized scene renders that we constructed to be similar to their own typical scene drawings or scene renders that were similar to other participants’ drawings. In line with previously reported data (19), scene renders that were typical for individual participants were categorized more accurately by these participants. Crucially, multivariate analyses on the EEG data allowed us to reveal how such effects emerge on the neural level. These analyses yielded two key results: First, perceptual representations of personally typical scenes emerging at 200ms of visual analysis are enhanced, compared to more atypical scenes. Second, personally typical scenes are represented in idiosyncratic ways, rendering the representations less faithful to the visual attributes of the scene.

## Results

### Personally typical scenes are better categorized

We used drawing as a behavioral readout of individual participants’ conceptions of typical everyday scenes (19). In a first drawing session, participants were asked to draw typical scenes from six categories (bathroom, bedroom, café, kitchen, living room, office). To control the potential effect of memory on the following task, participants also copied photographs from the six categories. To reduce the influence of individual differences in drawing ability and style, we transformed these drawings into standardized 3D renders (Fig. 1A), which were used as experimental stimuli in the second experimental session. Here, we tested if participants are more accurate in categorizing scenes that are similar to their typical drawings (i.e., personally typical scenes). To this end, participants completed a difficult 6-way categorization task (Fig. 1A). Participants viewed personally typical renders, based on their own typical drawings (“own” condition), renders typical for other participants, based on other participants’ typical drawings (“other” condition), and renders based on copied photographs (“control” condition). Behavioral categorization results showed higher categorization accuracy for the own condition (Fig. 1B), compared to both the other (t(50)=3.70, p<0.001; Fig.1B) and control (t(50)=4.25, p<0.001; Fig.1B) conditions. Response times were not different between conditions (F(2,49)=0.18, p=0.83). These results indicate that scene perception is more accurate for scenes that are typical for individual participant.

**Figure 1.**
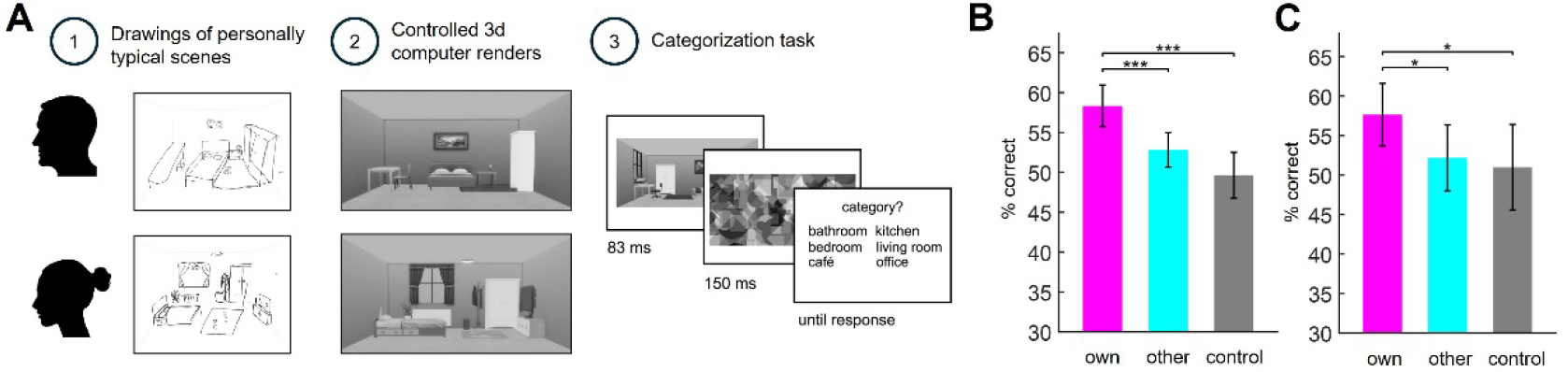
(A) Participants drew typical versions of six scene categories and copied scene photos from the same categories. To control for drawing ability differences across individuals, we converted drawings to 3D renders based on each participant’s own typical drawings, other participants’ typical drawings and the copied scenes. During the following categorization task, participants categorized briefly presented renders into six scene categories. (B) Behavioral categorization accuracy was higher for renders based on participants’ own drawings (own condition) than those based on other participants’ drawings (other condition) or copies (control condition). (C) We also separately analyzed the data from the 16 participants not yet reported before (19), the results were independently replicated. Error bars represent SEM. * indicates p < .05, ***indicates p < .001.

As these behavioral data were partly reported before – our previous behavioral report features 30 out of the 46 datasets (19) – we further tested whether the effect was independently replicated in the 16 participants not included in the previous study. The results from this new sample indeed fully replicated the effect, with significantly higher categorization accuracy in the own condition, compared to both the other (t(15)=2.33, p=0.034) and control (t(15)=2.26, p=0.039) conditions (Fig. 1C).

### Personally typical scenes evoke enhanced neural representations

To reveal how the personal typicality of a scene affects its representation in the brain, we recorded participants’ EEG signals while they performed the categorization task. We then conducted a time-resolved decoding analysis (20) on the EEG data to discriminate between the six scene categories (Fig. 2A), from -200 to 800 ms relative to scene onset, separately for the own, other, and control conditions. The resulting decoding performance over time yielded an estimate of the representational quality across conditions. Decoding accuracy rapidly increased for all conditions, starting from around 70 ms, and peaked around 200 ms (Fig. 2B). Critically, we found stronger decoding for the own condition, compared to both the other condition (from 175 ms to 225 ms), and the control condition (from 175 ms to 225 ms and at 275 ms). This result demonstrates an enhanced neural representation of personally typical scenes. The timing of this effect further suggests scene typicality on the individual level facilitates neural processing during perceptual analysis (21-24).

**Figure 2.**
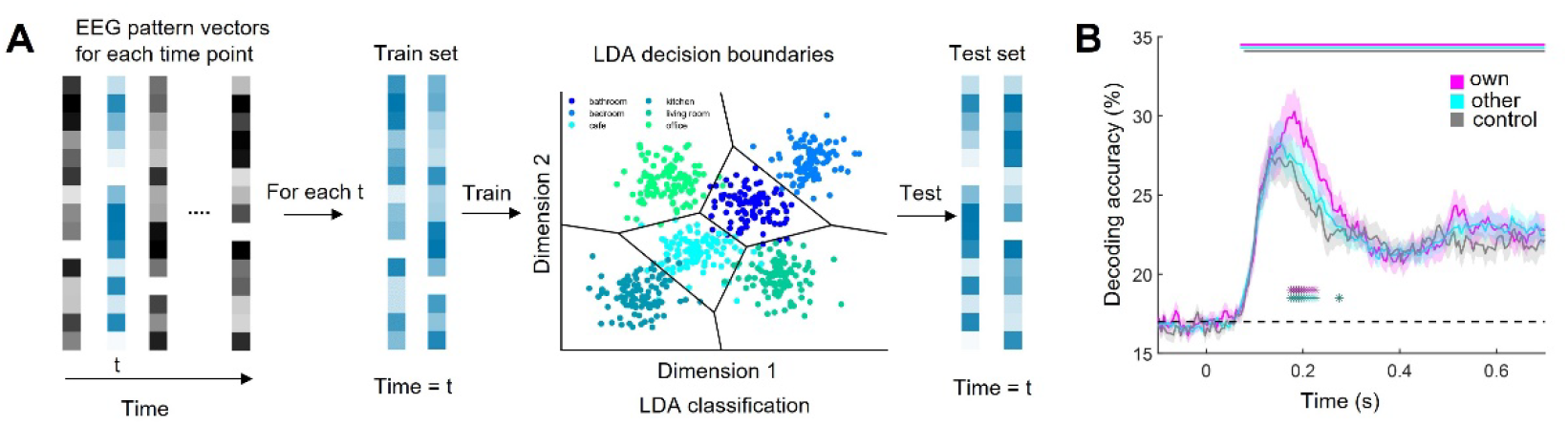
(A) To temporally track scene representations, we trained linear classifiers on the EEG activity pattern vectors at each time point. Classifiers were trained and tested on discriminating the six scene categories, using a 10-fold cross validation framework. (B) Between 175 ms and 225 ms after scene onset, decoding accuracy was higher in the own condition than in the other condition (indicated by dark purple significance markers) and the control condition (indicated by dark green significance markers). Error margins represent SEM. * indicates p < .05 (corrected for multiple comparisons).

### Personally typical scenes yield idiosyncratic neural representations

The EEG decoding results show that scenes that are typical for individual participants evoke enhanced cortical representations. These enhanced representations, however, could originate from two different underlying processes. On the one hand, these enhanced representations could stem from an enhanced representation of visual features, caused by sharper neural tuning to features that are prevalent in personally typical scenes. On the other hand, the enhanced neural representations for personally typical scenes could stem from these scenes more readily evoking higher-level representations that abstract away from visual features, for instance by more readily activating representations of categorical prototypes (25). Such higher-level visual representations could yield highly category-specific response patterns that boost decoder performance.

To arbitrate between these possibilities, we employed a deep neural network (DNN) model trained on scene categorization, which we used to quantify visual features extracted at different levels of the visual hierarchy. By quantifying how well the features extracted from the DNN predict brain activations in the EEG, we could test whether the enhanced neural representations of personally typical scenes are accompanied by a more or less pronounced representation of visual features. We first extracted activations from 12 layers along a googlenet (26) DNN trained on the Places365 dataset (27), for all scenes in the own, other and control conditions, separately for each participant. We then computed the pairwise similarity between these activations by correlating the activation vectors for each pair of scenes. This yielded a representational dissimilarity matrix (RDM) for each layer and condition. For the EEG, we first constructed RDMs based on the pair-wise decoding analysis and then averaged RDMs across the time window in which we found a difference between the own and other conditions (175 ms to 225 ms). To quantify how well the representation in the time window of interest are predicted by the visual feature organization in the DNN, we correlated the DNN RDM at each layer with the EEG RDM, separately for the three conditions (Fig. 3A). The results showed that representations of visual features in early and intermediate DNN layers similarly predict neural representations across three conditions (Fig. 3B). Critically, in the last two DNN layers, neural representations in the own condition were predicted less well by the DNN features, compared to both the other (inception5b layer: t(45)=2.71, p_FDR_=0.041; fully connected layer: t(45)=2.74, p_FDR_=0.018) and control (inception5b layer: t(45)=3.01, p_FDR_=0.041; fully connected layer: t(45)=3.50, p_FDR_=0.012) conditions (Fig. 3B). This shows that scenes that are typical for individual participants evoke neural representations that are less faithful to higher-level visual features extracted by the DNN. This supports the view that personally typical scenes more readily evoke categorical representations that abstract away from high-level visual properties. Interestingly, for the own condition, the correspondence between the DNN features and the neural representations gradually declined along the DNN’s feature hierarchy: When fitted a linear model on each participant’s data, we found a significantly negative slope across participants (t(45) = 3.05, p = 0.0038), which was absent in the other (t(45)=0.63, p=0.53) and control conditions (t(45)=0.47, p=0.64). This suggests that representations of personally typical scenes get progressively decoupled from visual features, compared to less typical scenes.

**Figure 3.**
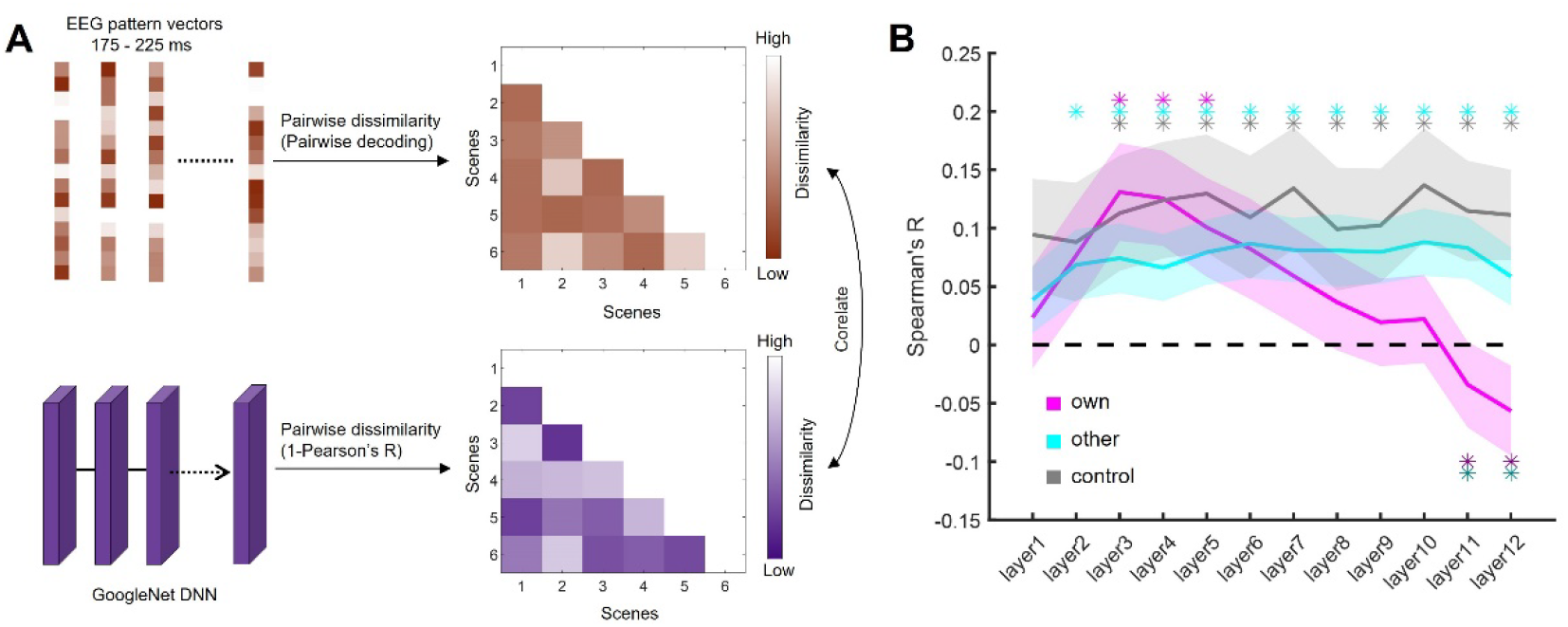
(A) To compare visual feature representations between a DNN and the brain, we first constructed EEG RDMs by computing pairwise decoding accuracies for all combinations of scenes, separately for the own, other, and control conditions. We then averaged RDMs for the time period showing a difference between the own and the other/control conditions (175 ms to 225 ms). Next, we constructed DNN RDMs by quantifying the pairwise dissimilarity of response vectors across the 12 layers of a GoogleNet trained on the Places365 dataset. Finally, we correlated the EEG and DNN RDMs for each condition, separately for each layer. (B) In the last two DNN layers, neural representations in the own condition were predicted less well by the DNN features, compared to both the other condition (indicated by dark purple significance markers) and the control condition (indicated by dark green significance markers). Error margins represent SEM. * indicates p < .05 (corrected for multiple comparisons).

## Discussion

Our findings demonstrate that the perception and neural representation of a scene vary as a function of whether the scene is typical for an individual participant. First, scenes that are more typical for individual participants are categorized more accurately, in line with previous results (19). Second, this enhanced categorization of personally typical scenes is accompanied by an enhanced cortical representation, emerging during visual analysis at around 200 ms post-stimulus. Finally, these enhanced representations are more idiosyncratic, that is, more detached from high-level visual features of the images.

These results are parsimoniously explained by predictive processing theories (28-30), which posit that perception is mediated by a convergence of feedforward sensory analysis and feedback in the form of predictions derived from internal world models. On this view, individual people –based on their individual lifetime experience with the world– form idiosyncratic internal models of what the world should look like. These individually specific internal models in turn yield individual differences in perception and neural representations. The enhanced neural representations for typical –and hence more predictable– scenes observed here are in line with a sharpening of cortical representations for predictable visual inputs (31, 32): Personally typical scenes may yield sparser neural codes that facilitate the readout of information.

Our results also reveal how representational idiosyncrasies emerge across the visual feature processing hierarchy. The DNN analysis shows that neural representations of personally typical and atypical scenes are similarly predicted by representations of low- and mid-level features in the DNN, suggesting that basic levels of visual processing do not reflect whether or not a scene taps into the internal model of individual observers (though one caveat here is that we purposefully matched the scenes in their low-level features). By contrast, representations are differently predicted by high-level features coding differs as a function of individual scene typicality (which is consistent with previous modelling of behavioral responses; 19): High-level visual features are coded less faithfully when the scenes are typical for individual participants. It furthermore suggests that personally typical scenes are represented in more idiosyncratic ways that are not captured by a DNN’s feature hierarchy. Neural processes that elude the modelling capacity of currently used DNN models include the activation of individually specific category prototypes (25, 33) or the rapid recruitment of semantic representations that bridge perception and memory (34). Such processes are likely to differ from individual to individual are thus not captured by DNN models trained on a single dataset. Alternatively, personally typical scenes may yield stronger recurrent connectivity that shapes more complex representations that are not captured by feedforward-only DNNs (35).

While our results highlight the enhancement of representations for personally typical scenes, they cannot directly elucidate the mechanisms through which these enhanced representations emerge from the interplay between feedforward and feedback dynamics in the cortex. To make progress, future research could combine EEG and fMRI recordings to chart the spatiotemporal dynamics evoked by personally typical and atypical scenes. Such studies could arbitrate between a purely feedforward emergence of sharpened representations for typical scenes and a refinement of these representations through cortical feedback from higher-level cortex or memory systems (36, 37). Future studies could also use DNN architectures that explicitly mimic feedback connectivity in cortex to disentangle feedforward and feedback information flows during the emergence of idiosyncratic representations in the visual system (35). For capturing the idiosyncratic nature of visual processing more comprehensively, these models could further be enriched with personalized training regimes or explicit participant-specific priors, which would push the limits of predictive power on the individual-participant level.

In sum, our research reveals idiosyncrasies in the way scenes are perceived and represented as a function of how typical they are to individual participants. Critically, we demonstrate that representations of personally typical scenes are enhanced and more idiosyncratic. These findings highlight that a comprehensive understanding of visual representations in human cortex requires researchers to take individual differences into account. With our drawing-based approach, we provide a straightforward method for predicting such differences.

## Materials and Methods

### Participants

Fifty-one participants (23.3±2.9 years, 15/36 male/female) completed the experiment. One additional participant completed the drawing session but did not return for the EEG session. Thirty-five participants were tested at Freie Universität Berlin, including EEG recordings for 30 participants (23.0±2.2 years, 8/22 male/female). Behavioral data for these participants has been reported before (19). Another 16 participants (23.9±4.3 years, 7/9 male/female) were tested at Justus-Liebig-Universität Gießen, using an identical setup. In total, 51 participants completed categorization task, including EEG recording for 46 participants (23.3±3.1 years, 15/31 male/female). The sample size for the EEG study is comparable to recent multivariate EEG studies conducted in the lab (36, 38, 39). Procedures were approved by the ethics committee of the Department of Education and Psychology, Freie Universität Berlin and the ethics committee of the Justus-Liebig-Universität Gießen, respectively, and adhered to the Declaration of Helsinki.

### Drawing sessions

Participants provided their drawings on an Apple iPad Pro using an Apple Pencil. Drawings were created using the Sketchbook App. Here, participants were asked to draw typical versions of six scene categories (bathroom, bedroom, café, kitchen, living room and office). For each drawing, they were given 30 s to plan the drawings and then 4 min to execute it. A perspective grid was provided to guide the arrangement of objects across the scene. We instructed participants to draw the most typical, but not their own or most liked, rooms. We additionally asked them to copy a photograph for each of the scene categories, under the same time constraints. Full details on the drawing session can be found in our previous study (19).

### Scene renders

Given the variation in participants’ drawing ability and style, we created experimental stimuli that removed these differences. Specifically, we created 3d renders similar to participants’ drawings by placing objects into an empty room, using the SIMS4 builder toolkit (see Fig. 1A). In total, 318 renders were created: 312 renders were based on the typical drawings of 52 individual participants and 6 renders were based on the 6 control photographs (these control renders were identical for all participants).

### Categorization task

We used the Psychophysics Toolbox (40) for Matlab to set up the categorization task. Here, participants were asked to categorize the briefly presented scene renders into six categories (see Fig. 1A). On each trial, a render (7° horizontal visual angle) was presented for 83 ms, followed by a geometric pattern mask, presented for 150 ms. Participants viewed renders based on their own drawing of a typical scene (“own” condition), based on other participants’ drawings of typical scenes (“other” condition), and based on their copied scenes (“control” condition). We grouped participants into groups of 4, and each participant saw renders based on their own drawings, renders based on other 3 participants’ drawings, and the control renders. Each of these 30 stimuli was repeated 40 times, for a total of 1200 trials, presented in random order.

### EEG recording and preprocessing

We recorded EEG signals using an EASYCAP 64-electrode system and Brainvision actiCHamp amplifier. Electrodes were arranged according to the 10-10 system. The data was recorded with 1,000 Hz sample rate and filtered online between 0.03 and 100 Hz. All electrodes were referenced online to the Fz electrode and re-referenced offline to the averaged signal from all channels. We used FieldTrip (41) to process offline EEG data. The continuous EEG data were segmented to the epoch into trials ranging from 200 ms before stimulus onset to 800 ms after stimulus onset, and baseline corrected by subtracting the mean of the pre-stimulus interval for each trial and channel separately. Trials containing excessive noise were removed by visual inspection. Blinks and eye movement artifacts were removed using independent component analysis and visual inspection of the resulting components. The epoch data were down-sampled to 200 Hz.

### Time-resolved decoding analysis

To trace the temporal representation of scenes across time, we performed a time-resolved multivariate decoding analysis using CosMoMVPA (42). In this analysis, we decoded between the six scene categories, separately for the three conditions (i.e., own, other, and control). Specifically, we performed classification analysis from 200 ms before stimulus onset to 800 ms after onset, using on a sliding time window (50-ms width, 5-ms resolution). For each time window separately, we used linear discriminant analysis (LDA) classifiers with 10-fold cross validation. We first allocated the EEG data to 10 folds randomly. LDA classifiers were then trained on data from 9 folds and then tested on data from the left-out fold. Prior to classification, principal component analysis (PCA) was performed on all data from the training set, and the PCA solution (retaining 99% of the variance) was projected onto the testing set (43). The amount of data in the training set was always balanced across scene categories. Classification was done repeatedly until every fold was left out once and accuracies were averaged folds. For the other condition, we performed separate classification analyses for the renders from each of the other 3 participants, and accuracy was averaged across these three analyses.

### Representational similarity analysis

To better understand how neural representations in the own, other, and control conditions are predicted by visual image features, we related neural representations to a DNN model using representational similarity analysis (RSA). We first extracted neural RDMs separately for the EEG data and DNN model, and then tested how well the DNN RDMs predict the EEG RDMs. For the EEG data, representational dissimilarity matrices (RDMs) were constructed using the same classification routine as in the original decoding analysis (see above), but now computing pairwise decoding accuracies for all possible combinations of six categories, separately for each of the three conditions. As we were particularly interested in representations during the time points that showed a difference between the own and other (and own and control) conditions, we performed this analysis on all time points between 175 ms to 225 ms, and then averaged RDMs across these time points. For the DNN, we extracted activation vectors from 12 layers (cov1, cov2, inception3a, inception3b, inception4a, inception4b, inception4c, inception4d, inception4e, inception5a, inception5b, full-connected) along the hierarchy of googlenet (25) trained on scene categorization using the Places365 dataset (26), as used in our previous study (19). We extracted layer-wise features activations for each of the 6 scenes used in each condition. We then constructed layer-specific RDMs by quantifying the pairwise dissimilarity (1-Pearson’s R) of response patterns in each layer. Finally, we correlated (Spearman’s R) the EEG RDMs with the DNN RDMs, separately for each layer and each condition. We Fisher-transformed the correlation values prior to statistical analysis.

### Statistical analysis

For the behavioral data analysis, responses slower than 5s were discarded. Accuracies and response times were compared using t-tests. Only trials with correct responses were analyzed for the response times. For the decoding analysis, we used t-tests and threshold-free cluster enhancement (TFCE; 44) in CoSMoMVPA to test decoding accuracies against chance level and assess differences between conditions. Multiple comparison correction across time was based on a sign-permutation tests with null distribution created from 10,000 bootstrapping iterations. For the RSA, we used t-tests to compare correlations against chance level and compare difference between conditions. Multiple comparison correction across layers was performed using false-discovery-rate (FDR) correction.

## Data, Materials, and Software Availability

Data, materials and code for the study are available at https://osf.io/ctsxv/.

## Acknowledgments

G.W. is supported by a PhD stipend from the China Scholarship Council (CSC). R.M.C. is supported by the Deutsche Forschungsgemeinschaft (DFG; CI241/1-1, CI241/3-1, CI241/7-1) and by a European Research Council (ERC) starting grant (ERC-2018-STG 803370). D.K. is supported by the DFG (SFB/TRR135, project number 222641018; KA4683/5-1, project number 518483074), “The Adaptive Mind”, funded by the Excellence Program of the Hessian Ministry of Higher Education, Science, Research and Art, and an ERC Starting Grant (PEP, ERC-2022-STG 101076057). Views and opinions expressed are those of the authors only and do not necessarily reflect those of the European Union or the European Research Council. Neither the European Union nor the granting authority can be held responsible for them.

## Author contributions

**Gongting Wang**: Writing – original draft, review & editing, Validation, Software, Methodology, Investigation, Formal analysis, Data curation, Conceptualization. **Lixiang Chen**: Writing – review & editing, Methodology. **Radoslaw M. Cichy**: Writing – review & editing, Supervision, Resources, Project administration, Funding acquisition. **Daniel Kaiser**: Writing – review & editing, Writing – original draft, Visualization, Validation, Supervision, Software, Resources, Project administration, Methodology, Funding acquisition, Conceptualization.

## Competing interests

The authors declare no competing interest.

## Notes

https://osf.io/ctsxv/

### Competing Interest Statement

The authors have declared no competing interest.

